# Tip Growth Defective1 interacts with the cellulose synthase complex to regulate cellulose synthesis in Arabidopsis thaliana

**DOI:** 10.1101/2023.09.15.557907

**Authors:** Edwin R Lampugnani, Staffan Persson, Ghazanfar Abbas Khan

## Abstract

Plant cells possess robust and flexible cell walls composed primarily of cellulose, a polysaccharide that provides structural support and enables cell expansion. Cellulose is synthesised by the Cellulose Synthase A (CESA) catalytic subunits, which form cellulose synthase complexes (CSCs). While significant progress has been made in unravelling CSC function, the trafficking of CSCs and the involvement of post-translational modifications in cellulose synthesis remain poorly understood. In order to deepen our understanding of cellulose biosynthesis, this study utilised immunoprecipitation techniques with CESA6 as the bait protein to explore the CSC and its interactors. We successfully identified the core proteins of the CSC complex and discovered new potential interactors involved in CSC trafficking and the coordination of cell wall synthesis. Moreover, we identified TIP GROWTH DEFECTIVE 1 (TIP1) protein S-acyl transferases (PATs) as an interactor of the CSC complex. We confirmed the interaction between TIP1 and the CSC complex through multiple independent approaches. Further analysis revealed that *tip1* mutants exhibited stunted growth and reduced levels of crystalline cellulose in leaves. These findings suggest that TIP1 positively influences cellulose biosynthesis, potentially mediated by its role in the S-acylation of the CSC complex.

## Introduction

Plant cell walls are complex and constantly changing structures made up of polysaccharides like cellulose, hemicelluloses, callose, and pectins (1). Primary cell walls are formed around every cell during the initial stages of cell division and expansion during development, while secondary cell walls are produced in specialised cells after the expansion phase has ended (1). Cellulose, which is made up of β-(1→4)-D-glucan chains, is the most abundant and primary load-bearing polymer in cell walls. It acts as a framework for the addition of other wall components and is synthesised by cellulose synthase A (CESA) catalytic subunits. These subunits are organised into large multiprotein complexes known as cellulose synthase complexes (CSCs) (2). There are ten members of the AtCESA family in plants, which can be divided into two groups: primary wall CESAs (AtCESA1, AtCESA3, AtCESA6, and AtCESA6-like (AtCESA2, AtCESA5, and AtCESA9) and secondary wall CESAs (AtCESA4, AtCESA7, and AtCESA8) (2). In addition to the CESAs, the CSC comprises various ancillary proteins, including COMPANION OF CELLULOSE SYNTHASEs (CCs), CELLULOSE SYNTHASE INTERACTIVE (CSI), and KORRIGAN (KOR) (3). CSCs are assembled in the Golgi apparatus and then transported to the plasma membrane, where cellulose synthesis occurs (4). CSCs move along the plasma membrane at a speed of 200-350nm/min to synthesise cellulose, and their movement is thought to be driven by the catalytic activity of the CESAs (5). Various developmental and environmental factors regulate CSC activity to alter tissue growth (6). For example, CSC activity can be regulated through the phosphorylation of CESAs in response to light and hormonal signalling (7). Additionally, CESAs can be modified through S-acylation, which involves the addition of acyl groups such as palmitate or stearate to cysteine residues (8). This modification can affect protein structure or localisation (9). CESAs involved in primary cell wall synthesis, including CESA2, CESA3, and CESA5, along with all CESAs involved in secondary cell wall synthesis (CESA4, CESA7, and CESA8), undergo S-acylation (8). In addition to the CESA enzymes involved in primary and secondary cell wall synthesis, several interacting proteins of the Cellulose Synthase Complex (CSC) and proteins associated with CESA trafficking have also been found to undergo S-acylation (8).

S-acylation, also called palmitoylation, is a reversible post-translational modification involving the transfer of a fatty acid group, typically stearate or palmitate, to a cysteine residue within target proteins (10). This modification can significantly impact protein properties due to the highly hydrophobic nature of the acyl group. In proteins lacking transmembrane helices (TMHs), S-acylation leads to their localisation in the membrane. Conversely, in proteins that already possess TMHs, S-acylation can result in various effects, such as altering subcellular localisation, partitioning into membrane microdomains, and facilitating the formation of multi-protein complexes (11).

A family of protein acyl transferases (PATs), characterised by a catalytic domain containing a DHHC motif, is responsible for adding the acyl group. In Arabidopsis, a total of 24 distinct PATs have been identified. These PATs are found in various cellular compartments, including the Golgi apparatus, endoplasmic reticulum, and plasma membrane (12). Tip Growth Defective1 (AtTIP1/AtPAT24) is a well-studied PAT in plants that possess ankyrin repeats, a DHHC-cysteine rich domain (CRD), and a PATs catalytic signature domain (13). AtTIP1 shows higher expression during various developmental stages compared to other members of the AtPATs family, suggesting its involvement in a wide range of developmental processes (12). Disruptions in AtTIP1 have been linked to shortened and branched root hairs, defects in pollen tube growth, cell polarity, cell size control, and reduced cell growth, resulting in a shorter overall stature (13, 14). The maize enzyme ZmTIP1 can interact with and S-acylate the protein ZmCPK9, leading to increased root hair elongation by mediating the association of ZmCPK9 with the plasma membrane (15).

In this study, we performed immunoprecipitation using CESA6 as the bait protein to investigate the CSC complex. Besides identifying the core CSC proteins, we also discovered new interactors involved in CSC trafficking and the coordination of cell wall synthesis. Additionally, our findings confirmed the interaction between TIP1 and the CSC complex through three independent approaches. *tip1* mutants exhibited stunted growth and a decrease in crystalline cellulose content in leaves. These observations suggest a positive role for AtTIP1 in cellulose biosynthesis, potentially mediated by its involvement in the S-acylation of the CSC complex.

## Materials and Methods

### Plant material and growth conditions

The T-DNA insertion lines, SALK_020996 (*tip1-3*) and SALK_089971C (*tip1-4*), were obtained from the Arabidopsis Biological Resource Center (ABRC). YFP-CESA6 lines were described previously (5). The primers used to identify these homozygous insertion lines can be found in Supplementary Table 1. For immunoprecipitation experiments, plants were grown in standard media on vertical agar plates with half-strength MS Basal Salt mixture, 0.5% MES, 0.5% Suc, and 0.8% Difco granulated agar. Arabidopsis seeds were sterilized with chlorine gas for three hours, resuspended in 0.1% agarose, stratified at 4°C for two days, and then planted in a single row on 12-cm-square plates sealed with 3M Micropore tape. These plates were placed in racks in a growth chamber with a 16-hour light/8-hour dark cycle, 120 E m2 s1 of light intensity, a temperature of 23°C during the day and 19°C at night, and 60% humidity. For phenotyping and cellulose quantification, single plants were grown in pots with standard potting mix and placed in a growth room with a 16-hour light/8-hour dark cycle, 120 E m2 s1 of light intensity, a temperature of 23°C during the day and 19°C at night, and 60% humidity.

### Co-immunoprecipitation-Mass spectrometry

To identify potential protein interactors of CESA6, we performed co-immunoprecipitation (Co-IP) experiments using GFP-Trap A beads. First, we ground and homogenized 1 g of tissue from plants expressing YFP-CESA6 or LTi6b-GFP as a control in Co-IP extraction buffer (100 mM Tris-HCl, pH 7.5, 150 mM NaCl, 1 mM EDTA, 10 mM MgCl2, 10% glycerol, 0.2% Nonidet P-40, 1 mM PMSF, 5 mM DTT). The protein extracts were incubated on ice for 30 min and then centrifuged to remove any debris. The clear supernatants were transferred to new tubes and the total protein concentrations were measured using the Bradford Assay. Next, we mixed 5 mg of total protein extracts with 200μl of GFP-Trap A beads and incubated the mixture overnight at 4°C with end to end rocking. After incubation, we washed the beads with TEN buffer (10 mM Tris-HCl, pH 7.5, 150 mM NaCl, 0.5 mM EDTA) and then with higher stringency TEN buffer (10 mM Tris-HCl, pH 7.5, 500 mM NaCl, 0.5 mM EDTA). The proteins bound to the beads were resuspended in 2 ml of TEN buffer. We performed on bead digestion with Trypsin followed by reduction with DTT and alkylation with Iodoacetamide. Five microliters of the protein digests prepared from the on-bead samples were analyzed using LC-MS/MS on a Q-executive mass spectrometer (Thermo Fisher Scientific). Data analysis was performed using the Proteome Discoverer 2.4 software (Thermo Scientific). A database search was performed using Proteome Discoverer version 2.4 with the SequestHT search engine and an *Arabidopsis thaliana* database (Tair10). The search parameters included Trypsin as the cleavage enzyme, precursor and fragment mass tolerances of 10 ppm and 0.6 Da respectively, and a maximum of 2 missed cleavages. Fixed modifications included carbamidomethyl cysteine, and variable modifications included oxidation of methionine and acetylation of the protein N-terminus. Protein and peptide groups were set to a maximum false discovery rate (FDR) of <0.01 as determined by the Percolator algorithm The mass spectrometry proteomics data have been deposited to the ProteomeXchange Consortium via the PRIDE (16) partner repository with the dataset identifier PXD045340.

### The split-ubiquitin membrane-based yeast two-hybrid system

To investigate the interactions between TIP1 and CSC proteins, we employed the split-ubiquitin membrane yeast two-hybrid assay as previously described (17). In our experimental setup, we cloned the TIP1 cDNA upstream of the C-terminal half of ubiquitin (Cub) and the synthetic transcription factor LexA-VP16 within the pBT3-SUC bait vector. For CESA3, CESA6, and CC1, their respective cDNAs were individually fused to the mutated N-terminal half of ubiquitin (NubG) within the pPR3-N prey vector. Co-transformation of the bait/prey pairs was accomplished in the NMY51 yeast strain, and the transformed cells were plated on selective medium lacking leucine and tryptophan (-Leu/-Trp). The physical interactions between the bait and prey was discerned by the growth of colonies on selective medium deprived of adenine, leucine, tryptophan, and histidine (-Ade/-Leu/-Trp/-His), with the inclusion of 5mM 3-amino-1,2,4-triazole to enhance stringency. The bait pTSU2-APP and the prey pNubG-Fe65 were used as positive controls.

### Bimolecular fluorescence complementation

Coding sequences of TIP1, CESA1, CESA3, CESA6 and CC1 were amplified from cDNA using primers defined in Supplemental Table 1 and inserted into pURIL, pDOX, pAMON, or pSUR vectors using the Gibson assembly method. The pURIL and pDOX vectors were cut with KpnI and SfoI enzymes, while pAMON and pSUR were cut with BamHI and SfoI. TIP1 was inserted into pURIL and pDOX, while NKS1, CESA3, CESA6, and CC1 were inserted into pAMON and pSUR to generate C-terminal and N-terminal fusions respectively. The resulting constructs were introduced into Agrobacterium and combined with 35S::CFP-N7 and P19 strains, which were then introduced into *N. benthamiana* leaves via infiltration according to the method described by Zhang et al. (2016). Fluorescence of *N. benthamiana* leaves was observed 3 days after infiltration with combinations of BiFC constructs and Agrobacterium strains carrying 35S::CFP-N7 and P19. The observations were made using an inverted Nikon Ti-E microscope equipped with a CSU-W1 spinning disk head (Yokogawa) and a 100× oil-immersion objective (Apo TIRF, NA 1.49). The iXon Ultra 888 EM-CCD (Andor Technology) was used to detect fluorescence. The same imaging conditions were used for all BiFC combinations, with a 445-nm laser line used to excite CFP and a 515-nm laser line used to excite YFP. Emissions were detected using 470/40 and 535/30 band-pass filters, and z-stacks of images were collected with exposure times of 100 ms.

## Results

### Immunoprecipitation of CSC complex

To identify potential interacting proteins of the CSC complex, we employed a YFP-CESA6 line in pulldown experiments. Immunoprecipitation was conducted on 10-day-old seedlings cultivated in liquid half-MS media. Subsequently, mass spectrometry analysis was employed to identify potential interactors of the CSC complex. As a control, we utilised GFP-tagged LTI6B, a protein localised to the plasma membrane (18). Using a rigorous analysis approach, we established specific criteria for our study, requiring an adjusted p-value below 0.05, an increase in abundance ratio of greater than 2 and the detection of at least two unique peptides per protein. Applying these criteria, we successfully identified the core components of the primary wall CSC, namely CESA1, CESA3, CESA6, CC1 and CSI1 (Fig. 1A, Supplementary Table1).

**Fig. 1:**
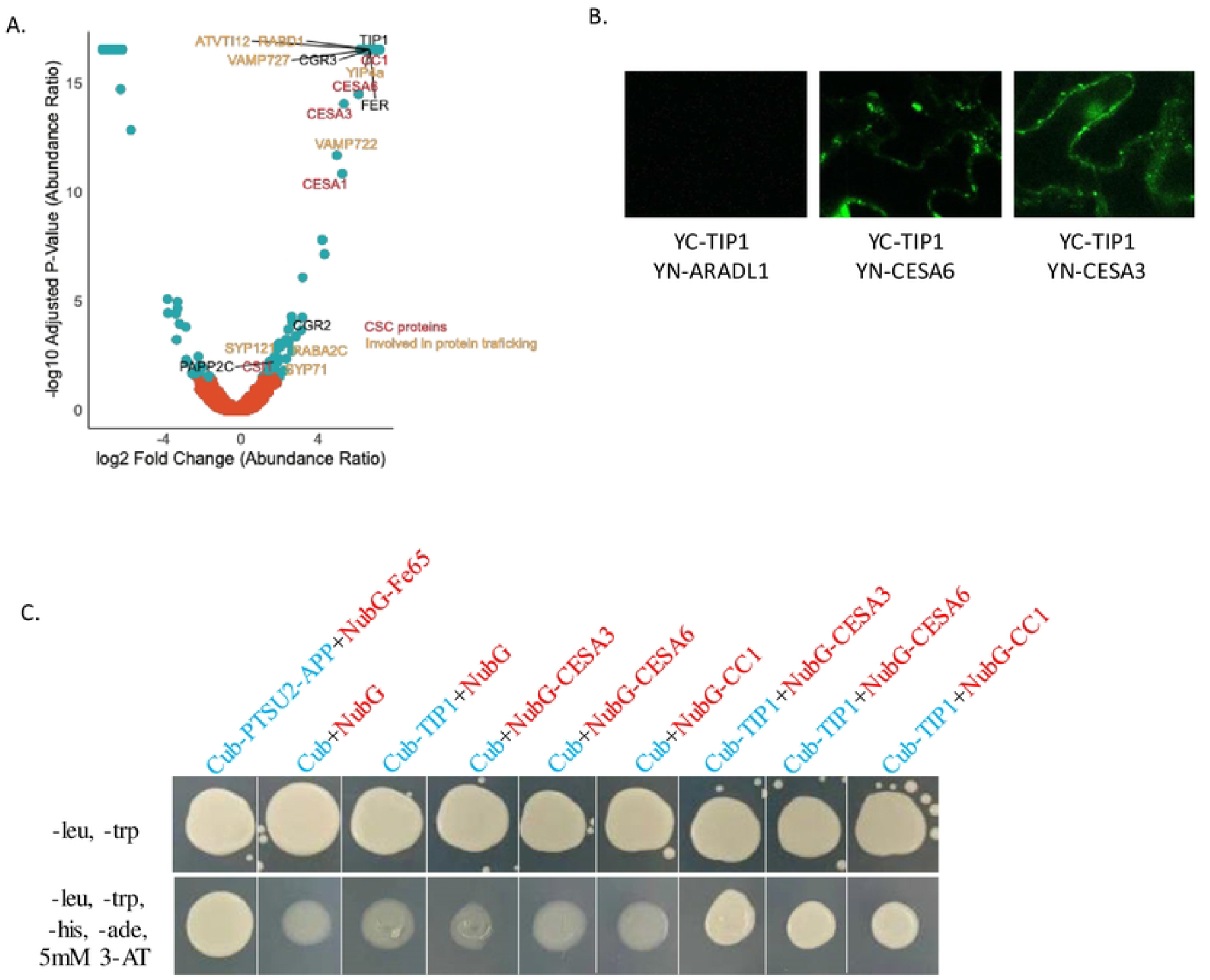
TIP1 interacts with the CSC complex. A. Volcano plot depicting proteins immunoprecipitated with the YFP-CESA6 bait in comparison to the Lti6b control. Proteins associated with the Cellulose Synthase Complex (CSC) are highlighted in green, while proteins involved in protein trafficking are highlighted in yellow. B. BiFC Assay Demonstrating Interactions between TIP1 and CESA1 and CESA3 Transiently Expressed in Epidermal Cells of *N. benthamiana* Leaves (Also se supports Figure 1.) N-terminal YN fusion of ARADL1 was used as controls and did not interact with YC fusions of TIP1. C. Split-ubiquitin-based membrane yeast two-hybrid assays were performed to examine the interactions between CESA1, CESA3, and CC1 with TIP1. In this experiment, TIP1 was utilised as the bait, and it was tagged at its C-terminus with a fusion of Cub (C-terminal fragment of ubiquitin). Conversely, CESA3, CESA6, and CC1 were tagged at their N-termini with NubG (a mutant of the N-terminal fragment of ubiquitin). The results of this assay demonstrated the interaction between TIP1 and CESA3, CESA6, as well as CC1, as evidenced by colony growth on the selection media. The bait pTSU2-APP and the prey pNubG-Fe65 were used as positive controls. Growth assays were conducted on selective plates lacking leucine, tryptophan, adenine, and histidine, supplemented with 5mM 3-amino-1,2,3-triazole (3-AT), as well as on control plates lacking leucine and tryptophan. Plate images were captured after four days of incubation at 30°C.

Furthermore, our study revealed novel putative interacting proteins that may be associated with the trafficking of the CSC to the plasma membrane. These proteins include SYNTAXIN OF PLANTS 121 and 71 (SYP121 and 71), VESICAL TRANSPORT V-SNARE 12 (VTI12), and two vesicle-associated membrane proteins (VAMP), namely VAMP722 and VAMP727, along with two RAB GTPASE homologues, RABD1 and RABA2C (Fig. 1A, Supplementary Table1).

In addition to these findings, we have also identified two proteins, COTTON GOLGI-RELATED 2 (CGR2) and CGR3, that are associated with pectin methylesterification (18). Furthermore, we detected a receptor-like kinase FERONIA (FER) and PHYTOCHROME-ASSOCIATED PROTEIN PHOSPHATASE TYPE 2C as potential interactors of CSC, which may be involved in regulating the phosphorylation status of CSC proteins (Fig. 1A, Supplementary Table1).

The co-immunoprecipitation experiments revealed potential additional interactors that participate in various processes, such as the trafficking of membrane proteins, coordination of cell wall synthesis, and post-translational modifications. The identification of such interactors holds significant importance in enhancing our understanding of the mechanisms underlying cellulose biosynthesis and cell wall organisation.

### TIP1 interacts with CSC proteins

TIP1 was highly abundant in YFP-CESA6 immunoprecipitation but not detected in the LTi6B control, providing evidence of its interaction with the CSC complex (Figure 1A, Supplementary table 1). To validate the interaction between TIP1 and CSC complex proteins, we employed bimolecular fluorescence complementation (BiFC) assays. In this approach, TIP1 was fused to one half of YFP, and CESA3 or CESA6 was fused to the other half. The BiFC results confirmed that TIP1 interacts with two CSC proteins, including CESA3 and CESA6 (Figure 1B). N-terminal YN or YC fusions of Gogi localised ARADL1 and MUR3 (19) were employed as controls, displaying interactions exclusively with themselves and not with any of the N-terminal YN or YC fusions of TIP1 or the CESAs (CESA1, 3, and 6). Notably, CESA1, 3, and 6 exhibited dimerization, as indicated by fluorescence signals when co-expressing YN or YC fusions of CESAs with YN or YC fusions of the CESAs. As a positive transformation control, the nuclear marker CFP-N7 (cyan) was consistently utilized in all experiments (Figure 1B, Supplemental figure 1). To further substantiate these findings, Split-ubiquitin-based membrane yeast two-hybrid assay was employed to examine the interaction between TIP1 and three different CSC complex proteins. For this experiment, TIP1 served as the bait and was tagged at its C terminus with a moiety consisting of Cub (C-terminal fragment of ubiquitin) fused to the transcription factor LexA-VP16. CESA3, CESA6 and CC1 were tagged at their N terminus with the NubG (a mutant of N-terminal fragment of ubiquitin). This assay confirmed that TIP1 can interact with CESA3, CESA6, and CC1, indicated by colony growth on the selection media, providing additional evidence for the interaction between TIP1 and the CSC complex proteins (Figure 1C). In summary, our findings establish TIP1 as a component of the CSC complex and suggest its potential involvement in cellulose synthesis.

### TIP1 mutants have reduced crystalline cellulose in the leaves

Two independent knockout mutants of TIP1 exhibited stunted growth, a characteristic trait observed in mutants with cellulose deficiency (Figure 2A). To assess the impact of TIP1 on cellulose production, we quantified cellulose levels in 21-day-old soil-grown rosettes, as this developmental stage displayed the most pronounced phenotype. Our analysis revealed a significant reduction in crystalline cellulose content in the leaves of the *tip1* mutants compared to the wild type (Figure 2B), indicating that TIP1 plays a role in regulating cellulose synthesis in Arabidopsis.

**Figure 2:**
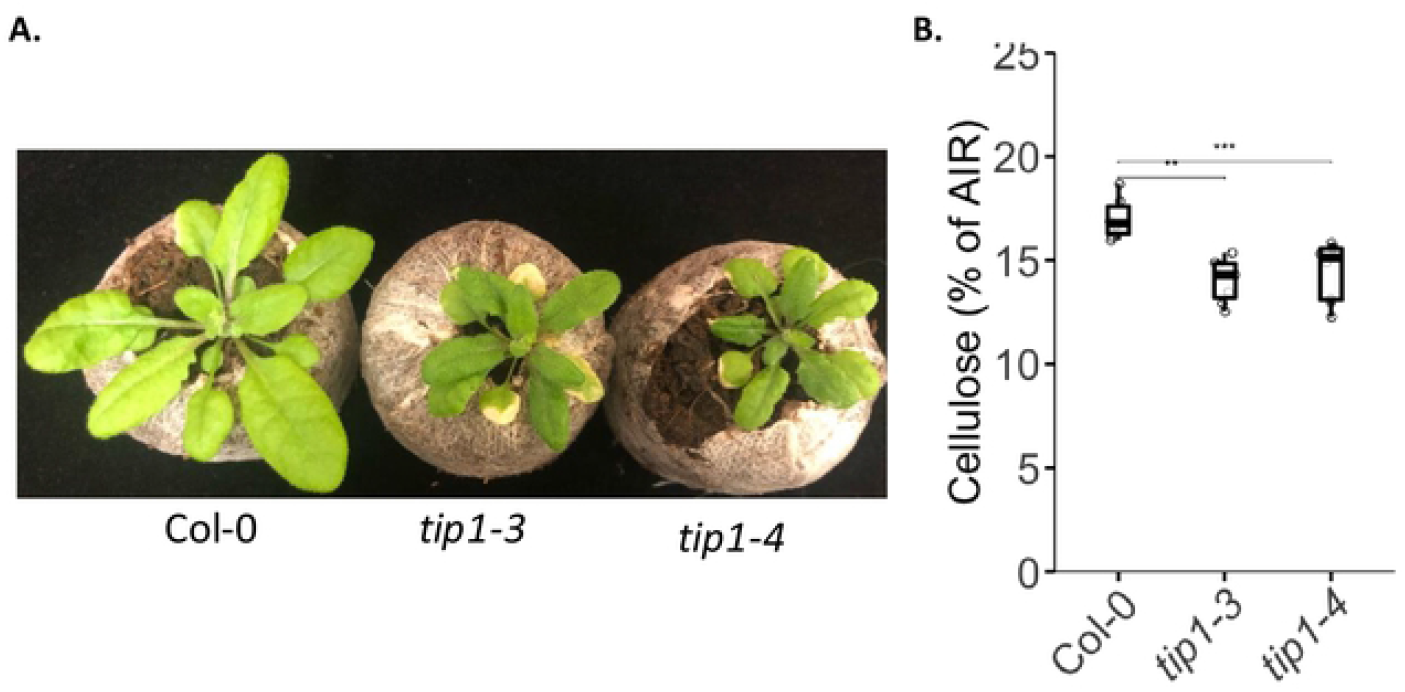
*tip1* mutants show stunted growth and reduced cellulose contents in leaves. A. Representative images illustrating stunted growth in Col-0, *tip1-3*, and *tip1-4* mutants. Seeds were directly sown in jiffy pots, and plants were grown under long-day conditions for 21 days prior to image capture. B. Cellulose quantification in the leaves of 21-day-old plants grown in jiffy pots. AIR: Alcohol insoluble residue. Data were obtained from five biological replicates of plants grown for 21 days and collected from three independent experiments. Asterisks indicate significant differences (Student’s t test, *P < 0.05 and **P < 0.01).

## Discussion

In our study, we utilised YFP-CESA6 as a bait protein for immunoprecipitation to uncover members of the CSC complex. This approach successfully identified CESA1, CESA3, CESA6, CC1, and CSI1, as part of the CSC complex. However, it is important to note that our analysis did not detect certain ancillary proteins of the CSC complex, including KORRIGAN. This limitation could be attributed to the choice of CESA6 as the bait protein or the possibility that these ancillary proteins exhibit weaker interactions, which may not have withstood the stringent conditions employed in our methodology.

In addition to the core CSC proteins, we have identified other interactors, specifically SYP121, VTI12, VAMP722, and VAMP727, which are integral components of the SNARE (soluble N-ethylmaleimide-sensitive factor attachment protein receptors) complex (20). The co-immunoprecipitation observed between these SNARE complex proteins and the CSC complex highlights a possible role of the SNARE complex in facilitating the targeting of CSC to the plasma membrane. Notably, VAMP727 establishes complexes with SYP121, both at the plasma membrane and in late endosomes, implying its involvement in membrane fusion events within these cellular compartments (20). While other SNARE proteins like SYP61 have been associated with CSC-containing trans-Golgi network (TGN) vesicles, their specific functional significance in CSC trafficking remains unclear (21). The identification of potential interactors of the SNARE complex opens up the possibility of unravelling the CSC trafficking pathways. The SNARE proteins, being integral components of membrane fusion events, have the potential to play a role in directing the targeting and trafficking of the CSC complex. However, a comprehensive understanding of the precise role of SNARE proteins in CSC trafficking necessitates further investigation.

A GTPase Interacting Protein 4a (YIP4a) was also identified as a potential interactor of the CSC complex. YIP4a, which forms a TGN-localized complex with ECHIDNA (ECH), which is required for the secretion of cell wall polysaccharides (22). The plant cell wall comprises interconnected structural components that form a unified, multi-network architecture. Thus, it is plausible that the cellulose biosynthesis and secretion of different cell wall polymers are tightly coordinated. The potential interaction of YIP4a suggests a likely coordination between the secretion of other cell wall-related polysaccharides and cellulose synthase complex (CSC) assembly, producing a well-organized cell wall structure. Further investigations are necessary to elucidate the precise role of YIP4a in cellulose biosynthesis and the organization of the cell wall.

Additionally, we identified a receptor like kinase FER and a protein phosphatase, namely PAPP2C, as a potential interactors of the CSC complex. Notably, CSC proteins undergo extensive phosphorylation, which significantly influences plant development (23). Despite this, the specific kinases or phosphatases responsible for regulating the phosphorylation status of CSC remain elusive. The identification of PAPP2C as a potential interactor within the CSC complex holds the potential to offer insights into the regulatory mechanisms governing CSC phosphorylation.

Finally, in our study, we observed the immunoprecipitation of TIP1 with CESA6, and we confirmed its interaction with the CSC complex using two independent approaches: BifC and split ubiquitination assays. Furthermore, we found that *tip1* mutants exhibited a reduction in crystalline cellulose content in the leaves compared to wild-type plants. These findings suggest that AtTIP1 plays a positive role in cellulose biosynthesis, potentially through the S-acylation of CSC proteins. During our work on this project, TIP1 was shown to be involved in the S-acylation of CESA3 (24). Furthermore, disrupting S-acylation sites of the secondary cell wall CESA7 leads to reduced trafficking of CESA7 to the plasma membrane, resulting in decreased plant height and crystalline cellulose content (9). Similarly, our assay using *tip1* mutants showed a reduction in crystalline cellulose content, suggesting that TIP1-mediated S-acylation of primary cell wall CESAs may be associated with their plasma membrane trafficking and, consequently, cellulose biosynthesis.

Lipid microdomains, enriched with sphingolipids, have been implicated in cellulose synthesis (25). Mutations in a Golgi-localized glycosyltransferase, resulting in impaired glycosylinositol phosphorylceramide (GIPC) synthesis, have shown decreased cellulose biosynthesis (25). The precise mechanism by which GIPCs affect cellulose synthesis is still unclear, but they may interact with cellulose synthesis components or indirectly modify plasma membrane properties and interactions. Considering that S-acylation forms microdomains within the plasma membrane, alterations in S-acylation could also influence cellulose biosynthesis by affecting the velocity of the CSC complex. These findings highlight the potential involvement of S-acylation in cellulose synthesis and warrant further investigation into their role and impact on CSC trafficking and mobility within the plasma membrane.

## Acknowledgements

ERL would like to acknowledge Australian Academy of Science, Thomas Davies Research Grant, The Australia & Pacific Science Foundation grant and the University of Melbourne Botany Foundation grant. SP was funded by a Villum, two Novo Nordisk, and Danish National Research Foundation grants (25915, 19OC0056076, 20OC0060564, DNRF155, respectively). GAK was funded by the Swiss National Science Foundation Fellowship (P2LAP3-168408), an ARC DECRA Fellowship (DE210101200) and an ABC grant from LA Trobe University.

## Figure legends

**Supplementary figure 1:**
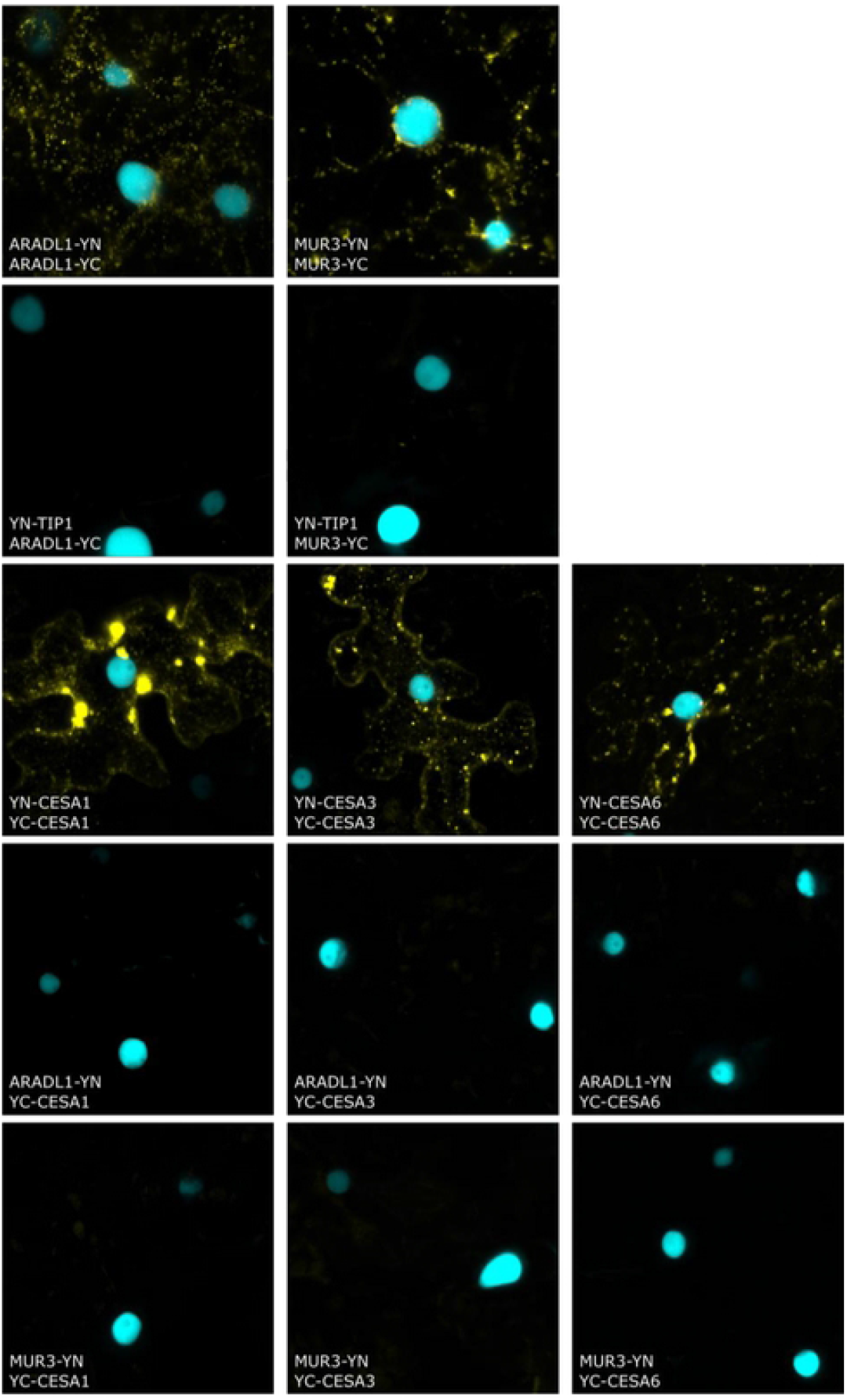
Controls for BiFC Assay. N-terminal YN or YC fusions of ARADL1 and MUR3 were used as controls and interacted only with themselves but not with any of the N-terminal YN or YC fusions of TIP1 or the CESAs (CESA1, 3 and 6). CESA1, 3 and 6 can dimerize as evidenced by fluorescence signal when co-expressing YN or YC fusions of CESAc with YN or YC fusions of the secondary wall CESAs The nuclear marker CFP-N7 (cyan) was used as a positive transformation control in all experiments.

